# Lactate Promotes Tunneling Nanotube Formation and Reduces Temozolomide Sensitivity in Glioblastoma Cells

**DOI:** 10.64898/2026.10.06.756927

**Authors:** Sambhavi Pattnaik, Subashree Anand, P Ganeshini, Amal Kanti Bera

## Abstract

Tunneling nanotubes (TNTs) mediate intercellular communication and contribute to cancer progression. Tumour cells produce excess lactate through increased glycolysis, but whether lactate regulates TNT formation has not been studied. Here, we show that lactate promotes TNT formation in U-87 MG glioblastoma (GBM) cells under both glucose-restricted and glucose-replete conditions, with a similar effect in MDA-MB-231 breast cancer cells. This effect depends on lactate uptake and metabolism rather than receptor-mediated signalling: inhibition of monocarboxylate transporter 1 (MCT1) or lactate dehydrogenase (LDH) reduced TNT formation, while exogenous lactate restored TNT formation during LDH inhibition. 3-chloro-5-hydroxybenzoic acid, an agonist of the plasma membrane-localized lactate receptor (GPR81/HCAR1), did not reproduce the effect of lactate. Lactate also increased intercellular mitochondrial transfer, attenuated temozolomide (TMZ)-induced growth suppression of GBM spheroids, and increased expression of the TNT-associated genes *CDC42*, *MYO10* and *FSCN1*. Inhibition of p300/CBP-mediated lactylation with A-485 reduced both protein lactylation and TNT formation, implicating lactylation in TNT biogenesis. Lactate increased basal and ATP-linked respiration that persisted during MCT1 inhibition and reduced on LDH inhibition, indicating that the effects of lactate on respiration and on TNT formation are at least partly separable. Together, these findings identify lactate as a regulator of TNT formation and link tumour metabolism to intercellular connectivity and TMZ response, highlighting lactate metabolism and lactylation as potential therapeutic targets.

**Highlights:**

- Lactate promotes tunneling nanotube (TNT) formation in glioblastoma and breast cancer cells.
- MCT1-dependent uptake and LDH-dependent metabolism, rather than GPR81 signalling, mediate lactate-induced TNT formation.
- p300-mediated protein lactylation contributes to lactate-induced TNT formation.
- Lactate enhances intercellular mitochondrial transfer and attenuates temozolomide-induced growth suppression of glioblastoma spheroids.

**Graphical abstract:** 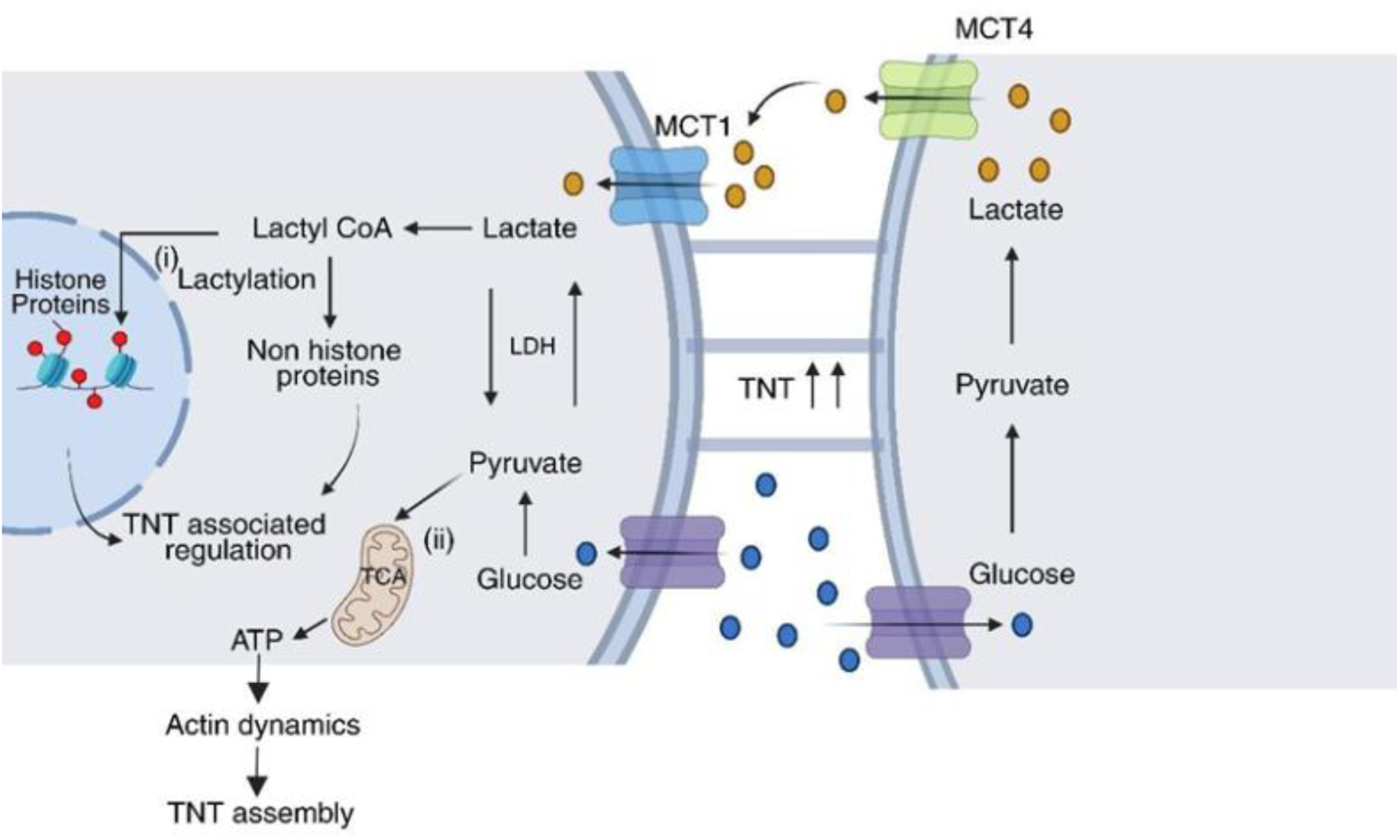

## Introduction

Glioblastoma (GBM) is the most aggressive and lethal form of primary brain tumour. Despite aggressive multimodal approach of maximal surgical resection followed by radiotherapy and temozolomide (TMZ) chemotherapy, median survival remains approximately 12–15 months. Recurrence is associated with the persistence of GBM stem cells and adaptation to the tumour microenvironment, which promote therapy resistance (Singh et al., 2025).

Tunneling nanotubes (TNTs) are thin, actin-based membrane structures that link distant cells, allowing the transfer of proteins, organelles and other cellular components (Belian & Zurzolo, 2026). First described in PC12 cells, TNTs have since been observed in diverse cell types and disease conditions (Rustom et al., 2004; Tiwari et al., 2021). In GBM, related structures known as tumour microtubes connect tumour cells into networks that promote invasion, growth and resistance to therapy (Osswald et al., 2015). TNT-mediated transfer of the DNA repair protein O6-methylguanine-DNA methyltransferase (MGMT) has also been implicated in TMZ resistance (Valdebenito et al., 2020).

The GBM microenvironment is hypoxic, nutrient-limited and metabolically reprogrammed, conditions that can influence TNT formation (Bayona et al., 2026; Valdebenito et al., 2021). Increased glycolysis and lactate production despite adequate oxygen availability are hallmarks of this metabolic state and are characteristic of the Warburg effect (Liberti & Locasale, 2016). Lactate accumulates in the tumour microenvironment and is increasingly recognised as both a signalling molecule and metabolic substrate that promotes tumour growth, invasion and therapy resistance (Pérez-Tomás & Pérez-Guillén, 2020). Because TNT formation is influenced by the metabolic and signalling state of cells (Belian & Zurzolo, 2026; Hekmatshoar et al., 2018), lactate may also regulate TNT formation. However, this possibility has not been directly tested in GBM.

Lactate crosses the plasma membrane primarily through proton-linked monocarboxylate transporters (MCT1–4). MCT1 is a bidirectional transporter that is highly expressed in GBM, correlates with poor outcome, and its inhibition sensitises GBM cells to TMZ (Behera et al., 2025; Miranda-Gonçalves et al., 2016). Lactate can also signal independently of cellular uptake through the G-protein-coupled receptor GPR81 (HCAR1), which has been linked to tumour progression and chemoresistance in several cancers (Ishihara et al., 2022; Longhitano et al., 2022). In addition, lactate promotes lysine lactylation of histone and non-histone proteins, providing a potential link between tumour metabolism and cellular function. This modification was first described by Zhang et al., who identified p300 as a lactyltransferase (Zhang et al., 2019). Lactylation has since been implicated in tumour progression and chemoresistance. In GBM, H3K9 lactylation (lysine at position 9 of histone H3) promotes TMZ resistance by activating LUC7L2, a regulator of RNA splicing, and suppressing MLH1, a key protein involved in DNA mismatch repair (Yue et al., 2024). H3K18 lactylation has also been associated with acquired TMZ resistance (Z. C. Wang et al., 2025). Whether lactylation contributes to the cytoskeletal remodelling required for TNT formation remains unknown.

Here, we show that extracellular lactate promotes TNT formation in GBM cells through MCT1-dependent uptake and LDH-dependent metabolism, independently of GPR81 signalling. This response is associated with p300-dependent lactylation, increased mitochondrial oxidative metabolism, enhanced TNT-mediated mitochondrial transfer and reduced sensitivity to TMZ. These findings identify a metabolic mechanism linking the lactate-rich GBM microenvironment to intercellular communication and chemoresistance.

## Results

### Lactate promotes TNT formation in GBM cells

We first examined whether extracellular lactate affects TNT formation in U-87 MG cells. Cells were labelled with CellMask™ Plasma Membrane Green to visualise and quantify TNTs. U-87 MG cells formed extensive TNT networks, and MitoTracker Red-labelled mitochondria were detected within some TNTs, consistent with their ability to transport organelles (Fig. 1A, B). We next cultured cells under different glucose and lactate conditions. Addition of 10 mM lactate increased TNT numbers under both glucose-restricted conditions (5.25 mM glucose versus 0.5 mM glucose + 10 mM lactate) and glucose-replete conditions (25 mM glucose versus 25 mM glucose + 10 mM lactate) (Fig. 1C-E). Under glucose-restricted conditions, approximately 160 TNTs were detected per 100 cells, increasing to nearly 300 TNTs per 100 cells following lactate addition. Under glucose-replete conditions, TNT numbers increased from approximately 250 to 340 per 100 cells after lactate treatment. Shorter TNTs are reported to be dynamic and exploratory, formed during the initiation phase as cells attempt to contact surrounding cells (Veranič et al., 2008). In this study, TNTs shorter than 75 µm were considered short TNTs, and lactate treatment increased the proportion of short TNTs (Fig. S1), consistent with enhanced TNT formation. Together, these results indicate that extracellular lactate promotes TNT biogenesis under both glucose-restricted and glucose-replete conditions.

**Fig. 1.**
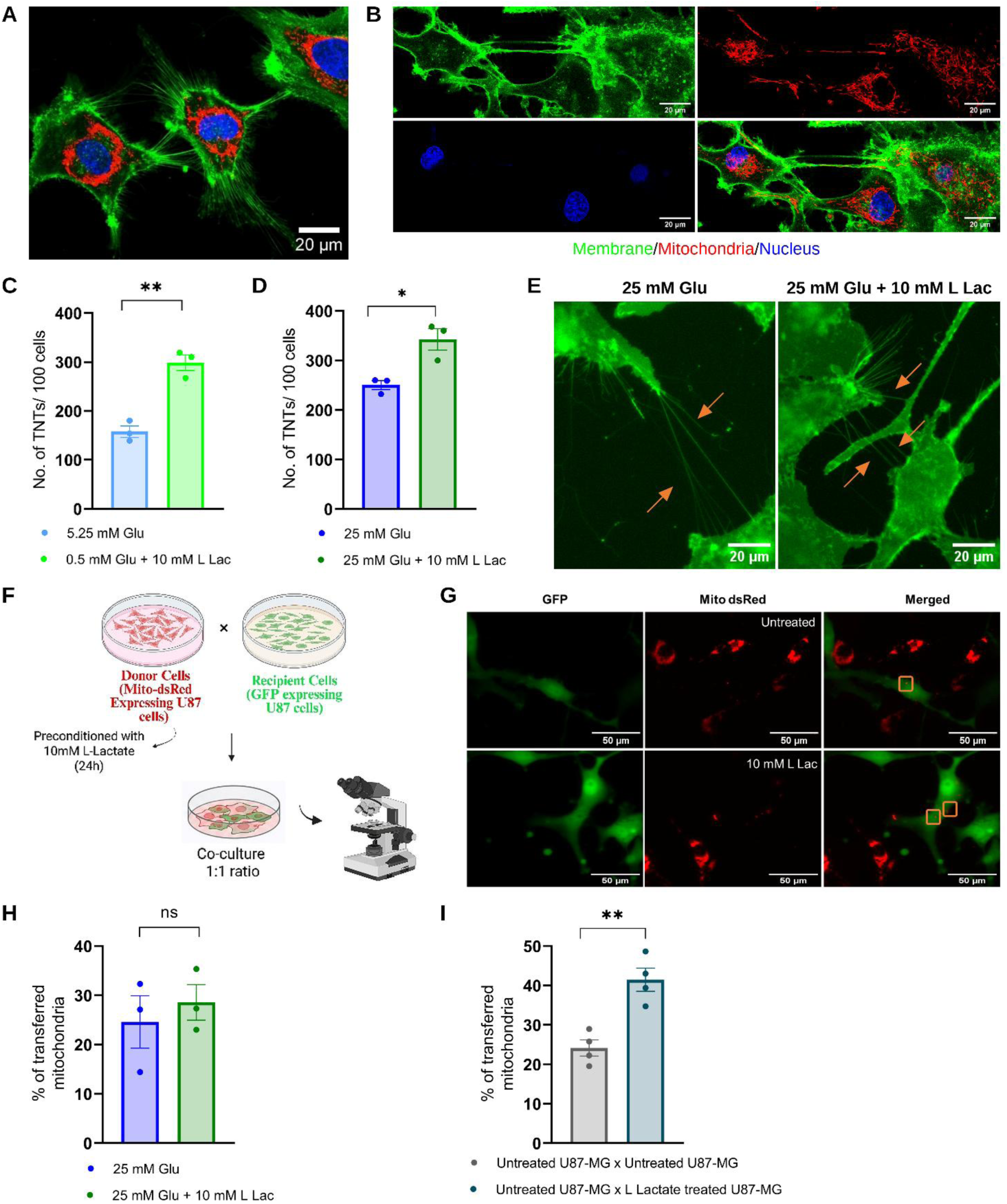
Lactate-driven TNT formation and mitochondrial transfer in GBM cells. **(A)** Representative confocal image of U-87 MG cells showing plasma membrane, mitochondria, and nuclei. **(B)** Representative confocal image showing mitochondria within a TNT. **(C, D)** Quantification of TNTs under glucose-restricted (C) and glucose-replete (D) conditions in the absence or presence of lactate. **(E)** Representative images of cells under glucose-replete conditions with or without lactate. **(F)** Schematic of the mitochondrial transfer assay. **(G)** Representative images showing mitochondrial transfer from donor to recipient cells. Boxes indicate DsRed-positive mitochondria in GFP-positive recipient cells. **(H)** Mitochondrial transfer (DsRed-positive puncta in GFP-positive recipient cells) when both donor and recipient cells were cultured with or without lactate. **(I)** Mitochondrial transfer from untreated or lactate-preconditioned donor cells to untreated recipient cells. Data are mean ± SEM from three independent experiments (C, D, H) or four independent experiments (I). *P < 0.05, **P < 0.01; unpaired t-test. ns, not significant. Scale bars, 20 µm (A, B, E) and 50 µm (G).

To determine whether this effect extends beyond GBM, we examined MDA-MB-231 breast cancer cells. Lactate increased TNT numbers under both low- and high-glucose conditions (Fig. S2), suggesting that lactate-induced TNT formation may represent a broader response to the tumour microenvironment. We next asked whether lactate-induced TNTs support mitochondrial transfer. Recipient cells stably expressing cytosolic GFP were co-cultured with donor cells expressing mitochondria-targeted DsRed (Fig. 1F, G). Mitochondrial transfer was quantified by measuring DsRed-positive mitochondrial puncta in recipient cells and normalising the values to recipient cell number. We tested two conditions: both donor and recipient cells were exposed to lactate during co-culture, or donor cells were preconditioned with lactate before co-culture with untreated recipient cells. When both donor and recipient cells were exposed to lactate, mitochondrial transfer showed an increased trend but did not reach statistical significance (Fig. 1H). However, preconditioning of donor cells with lactate significantly increased mitochondrial transfer (Fig. 1I). These findings indicate that lactate exposure, particularly donor-cell preconditioning, enhances mitochondrial transfer between GBM cells, consistent with increased TNT-mediated communication.

Because extracellular vesicles (EVs) can also contribute to mitochondria transfer, we examined whether EV-mediated transfer contributed to this effect. Cells containing DsRed-labelled mitochondria were incubated for 24 h in medium with or without lactate, after which the conditioned media were transferred to U-87 MG cells containing unlabelled mitochondria. After 16 h, DsRed-positive mitochondrial puncta were undetectable in recipient cells, with no apparent difference between cells treated with lactate-conditioned and control-conditioned media (Fig. S3). Thus, extracellular vesicle-mediated transfer appears to contribute minimally to the mitochondrial transfer observed under these conditions, supporting a predominant role for TNT-mediated cell-to-cell communication.

### Lactate uptake and intracellular metabolism, rather than receptor-mediated signalling, promote TNT formation

Lactate can act as a signalling molecule through the cell-surface receptor GPR81 (HCAR1) or as a metabolic substrate following transporter-mediated uptake (Li et al., 2022). To distinguish between these mechanisms, we first confirmed HCAR1 expression in U-87 MG cells by semi-quantitative PCR (Fig. S4). Treatment with the selective HCAR1 agonist 3-chloro-5-hydroxybenzoic acid (CHBA; 100 µM) for 24 h did not significantly increase TNT formation compared with control cells (Fig. 2A, B), suggesting that HCAR1 signalling does not mediate the effect of lactate. We next examined the role of MCT1 using the MCT1 inhibitor SR13800 (1 µM, 24 h). MCT1 inhibition significantly reduced TNT formation under both basal conditions (25 mM glucose) and in the presence of exogenous lactate (25 mM glucose + 10 mM L-lactate) (Fig. 2C-E). These findings indicate that MCT1-dependent lactate uptake contributes to TNT formation under both conditions.

**Fig. 2.**
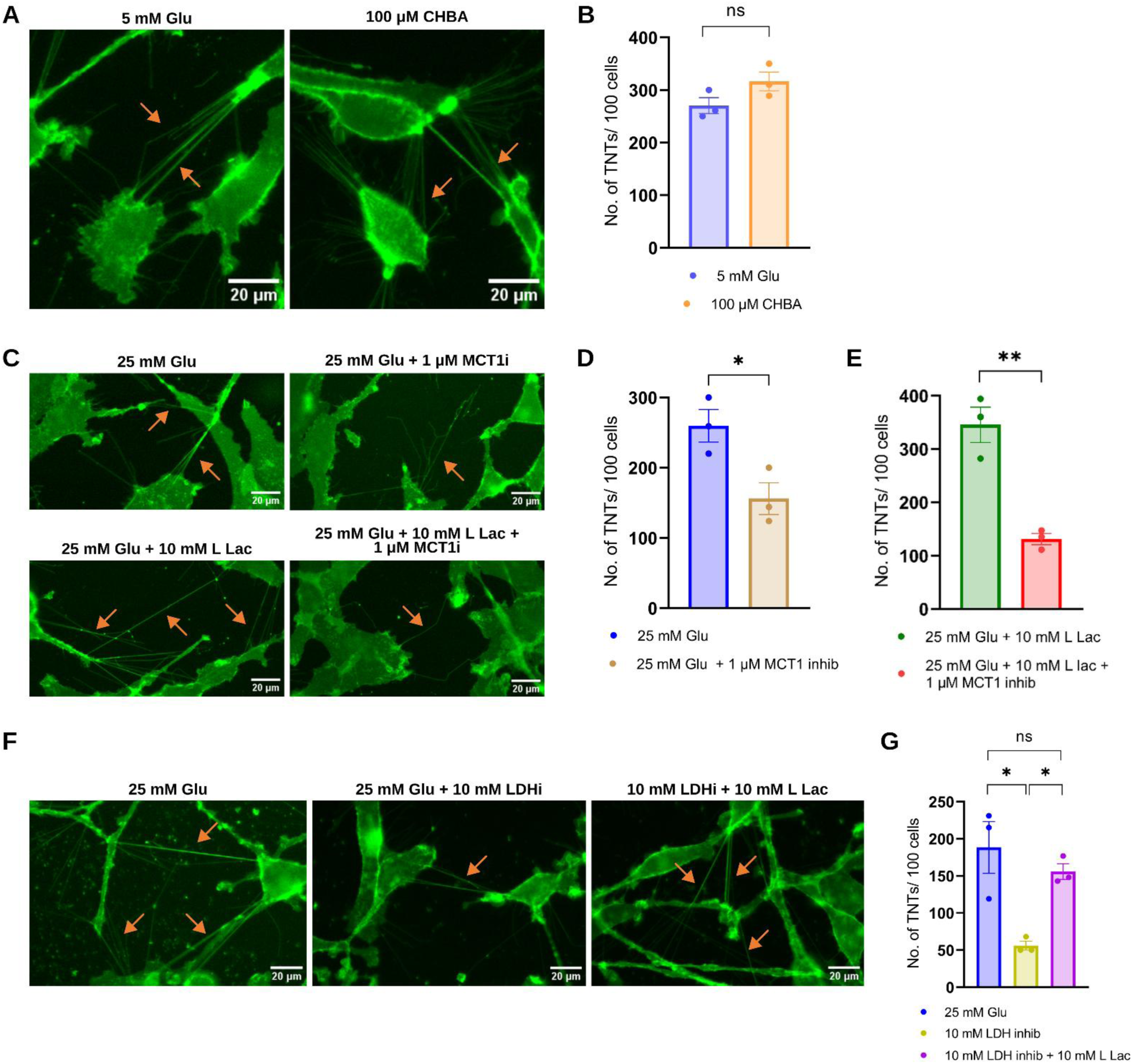
TNT formation is induced by lactate uptake and metabolism. **(A)** Representative images of U-87 MG cells treated with the HCAR1 agonist CHBA or control medium. **(B)** Quantification of TNTs under the conditions shown in (A). **(C)** Representative images of cells treated with lactate and/or the MCT1 inhibitor SR13800. **(D, E)** Quantification of TNTs in the absence or presence of SR13800 under basal (D) and lactate-supplemented (E) conditions. **(F)** Representative images of cells treated with the LDH inhibitor sodium oxamate with or without lactate. **(G)** Quantification of TNTs under the conditions shown in (F). Data are mean ± SEM from three independent experiments. *P < 0.05, **P < 0.01; one-way ANOVA (G) or unpaired t-test (B, D, E). ns, not significant. Scale bar, 20 µm.

GBM cells are highly glycolytic and produce substantial amounts of lactate from pyruvate through LDH (Guyon et al., 2022). We therefore tested whether endogenous lactate production contributes to TNT formation. Inhibition of LDH with sodium oxamate (10 mM, 24 h) significantly reduced TNT formation, and this reduction was reversed by exogenous lactate (10 mM) (Fig. 2F, G). These findings support a role for lactate metabolism in promoting TNT formation.

### Lactate reduces TMZ sensitivity in GBM spheroids and alters the expression of TNT-associated genes

TMZ can increase TNT formation, which contributes to drug resistance (Valdebenito et al., 2020). We therefore examined whether lactate-induced TNT formation affects the response of GBM cells to TMZ. U-87 MG-derived 3D spheroids were treated with 50 µM TMZ in the presence or absence of 10 mM lactate under low-glucose (5.25 mM glucose) or glucose-replete (25 mM glucose) conditions. Under low-glucose conditions, TMZ reduced the diameter of control spheroids approximately by 17%, compared with 5% in lactate-treated spheroids (Fig. 3A, B). Under glucose-replete conditions, the corresponding reductions were approximately 26% and 9%, respectively (Fig. 3C-E). Thus, lactate attenuated TMZ-induced growth suppression in GBM spheroids.

**Fig. 3.**
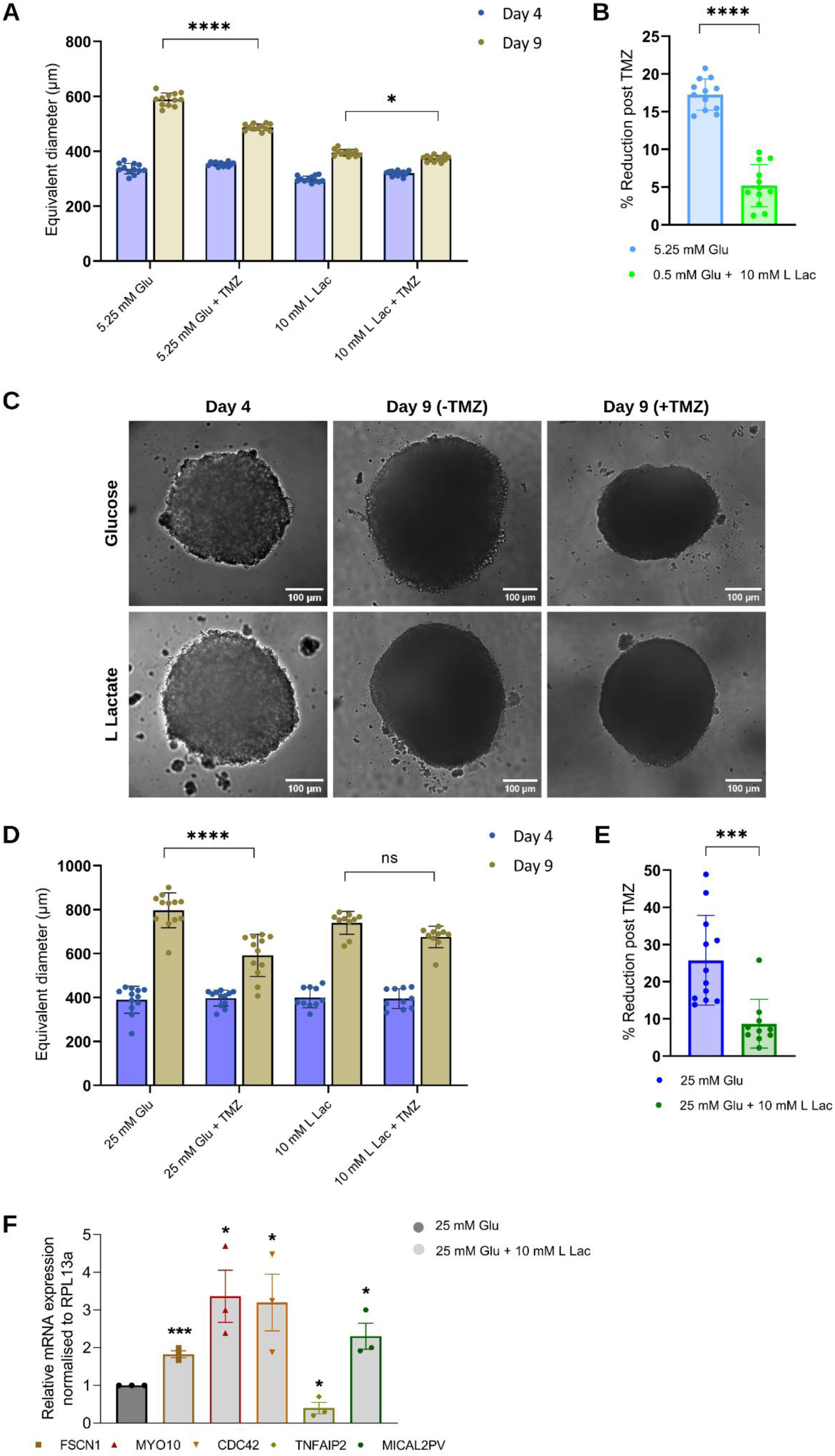
Lactate attenuates TMZ-induced growth suppression of GBM spheroids and alters the expression of TNT-associated genes. **(A)** Equivalent diameter of U-87 MG spheroids at days 4 and 9 under glucose-restricted conditions with or without lactate and TMZ. **(B)** Percentage reduction in spheroid diameter following TMZ treatment under the conditions shown in (A). **(C)** Representative phase-contrast images of spheroids before and after TMZ treatment under glucose-replete conditions with or without lactate. **(D)** Equivalent diameter of spheroids at days 4 and 9 under glucose-replete conditions with or without lactate and TMZ. **(E)** Percentage reduction in spheroid diameter following TMZ treatment under the conditions shown in (D). Spheroid data are mean ± SD (n = 10 - 12 spheroids per condition; each point represents one spheroid). **(F)** Relative mRNA expression of FSCN1, MYO10, CDC42, TNFAIP2, and MICAL2PV in lactate-treated cells compared with control cells. qPCR data are mean ± SEM from three independent experiments. *P < 0.05, ***P < 0.001, ****P < 0.0001; statistical tests are indicated in the respective panels. ns, not significant. Scale bar, 100 µm.

To examine molecular changes associated with lactate-induced TNT formation, we measured the expression of genes involved in TNT formation and actin remodelling. U-87 MG cells were cultured in 25 mM glucose with or without 10 mM lactate. Lactate significantly increased the expression of *CDC42*, *MYO10*, and *FSCN1* (Fig. 3F), which are associated with TNT formation and extension (Gousset et al., 2013; Hanna et al., 2017; Lou et al., 2012). Lactate also increased the expression of *MICAL2PV*, a negative regulator of TNT formation (F. Wang et al., 2021), and reduced the expression of *TNFAIP2*, a positive regulator of TNT biogenesis (Barutta et al., 2023) (Fig. 3F).

### Lactate promotes TNT formation through protein lactylation and enhances oxidative metabolism

Because lactate can promote protein lactylation (Zhang et al., 2019), we examined whether this modification contributes to TNT formation. Treatment with A-485 (0.5 µM, 24 h), an inhibitor of p300-mediated lactylation (Lasko et al., 2017), significantly reduced TNT numbers in lactate-treated cells (Fig. 4A, B) and decreased global protein lactylation (Fig. 4C, D). These findings support a role for p300-mediated lactylation in lactate-induced TNT formation downstream of lactate uptake.

**Fig. 4.**
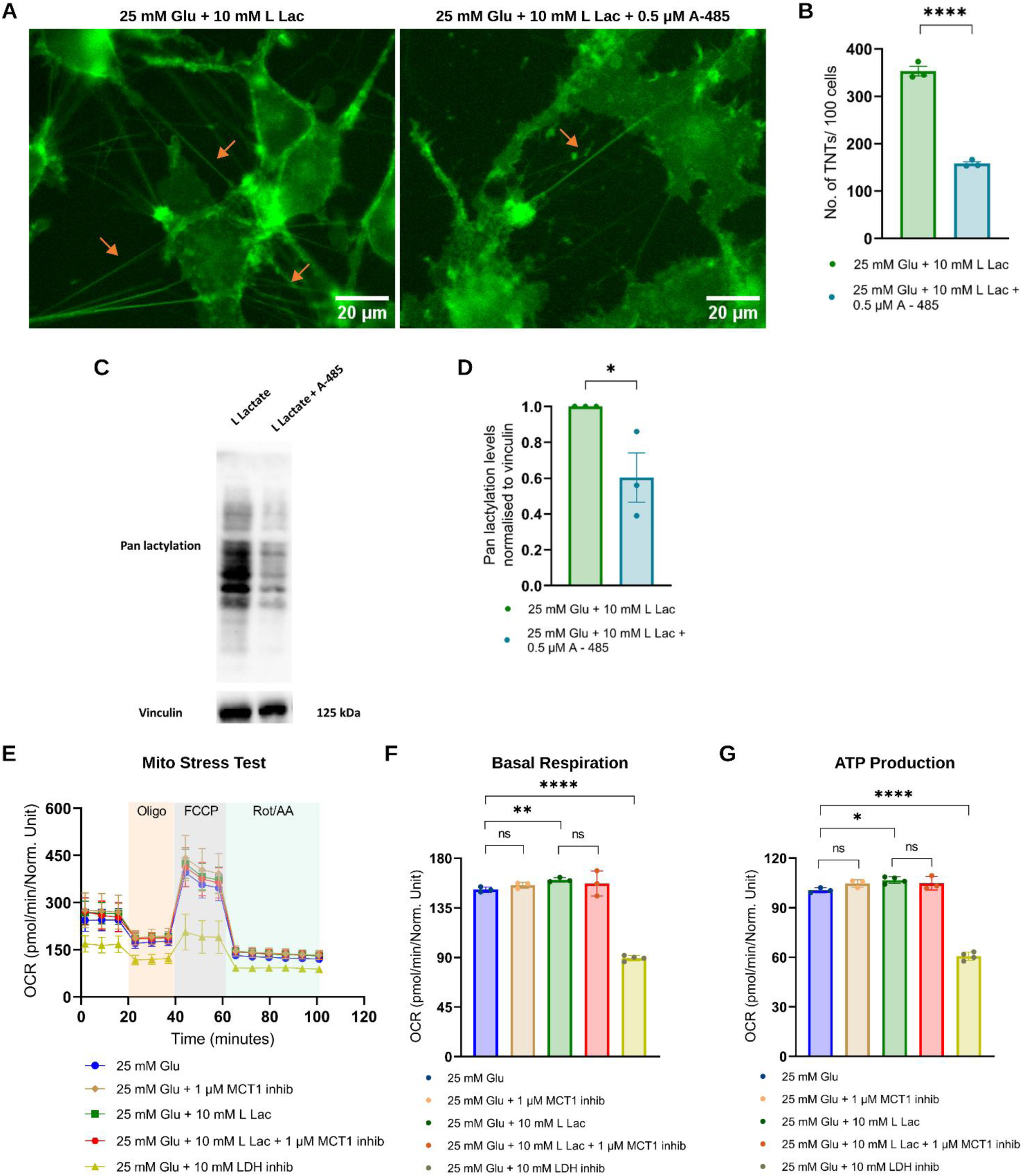
Lactate-driven protein lactylation and enhanced metabolism induce TNT formation. **(A)** Representative images of U-87 MG cells treated with lactate in the presence or absence of the p300/CBP inhibitor A-485. **(B)** Quantification of TNTs under the conditions shown in (A). **(C)** Representative western blot showing pan-lactylation. Data are mean ± SEM from 3 independent experiments. *P < 0.05, ***P < 0.001; unpaired t-test. Scale bar, 20 µm. **(D)** Quantification of pan-lactylation normalised to vinculin. **(E)** Mito Stress Test profiles showing oxygen consumption rate (OCR) under the indicated conditions. Oligomycin, FCCP, and rotenone/antimycin A injections are indicated. **(F, G)** Basal respiration (F) and ATP-linked respiration (G) derived from the Mito Stress Test. Data are mean ± SD from 3–4 independent measurements. *P < 0.05, **P < 0.01, ****P < 0.0001; unpaired t-test. ns, not significant.

TNT formation and mitochondrial transfer are energy-demanding processes. We therefore examined whether lactate alters mitochondrial oxidative metabolism using Seahorse extracellular flux analysis. Under glucose-replete conditions, 10 mM lactate significantly increased basal oxygen consumption rate (OCR) and ATP-linked respiration compared with glucose alone (Fig. 4E-G), indicating enhanced mitochondrial respiration. We next examined whether this response depends on the MCT1-LDH pathway. LDH inhibition significantly reduced basal and ATP-linked OCR, indicating that LDH-dependent conversion of lactate to pyruvate contributes to lactate-supported oxidative metabolism. In contrast, MCT1 inhibition did not significantly reduce the lactate-induced increase in OCR, although it significantly reduced TNT formation under the same conditions (Fig. 2E). Thus, the effects of lactate on mitochondrial respiration and TNT formation appear to be partly separable. LDH-dependent metabolism contributes to the oxidative response, whereas MCT1 activity has a more specific role in TNT formation.

## Discussion

We show that extracellular lactate promotes TNT formation in U-87 MG glioblastoma cells and enhances intercellular mitochondrial transfer and resistance to TMZ-induced growth suppression. Lactate increased TNT formation under both glucose-restricted and glucose-replete conditions and produced a similar effect in MDA-MB-231 breast cancer cells, suggesting that lactate-induced TNT formation may extend beyond GBM.

Our data indicate that lactate promotes TNT formation primarily through uptake and metabolism rather than GPR81 signalling. The GPR81 agonist CHBA did not reproduce the effect of lactate, whereas MCT1 inhibition reduced TNT formation under both basal and lactate-supplemented conditions. LDH inhibition produced a similar reduction that was reversed by exogenous lactate. The reduction in TNT formation following MCT1 inhibition under basal conditions (no exogenous lactate) may reflect the high glycolytic activity of GBM cells, in which lactate is produced and transported through MCT4 and can be re-imported through MCT1. Thus, MCT1 may represent an important control point for TNT formation.

A-485 reduced both global protein lactylation and TNT formation, supporting a role for p300-mediated lactylation in TNT biogenesis. Consistent with this finding, lactate increased the expression of *CDC42*, *MYO10*, and *FSCN1*, which are involved in actin remodelling and TNT formation (Brou & Zurzolo, 2026). Whether lactylation directly regulates these genes or instead affects the cytoskeletal machinery remains unclear. Future studies using locus-specific mapping or rescue experiments will be required to distinguish between these possibilities.

Lactate also increased *MICAL2PV* expression and reduced *TNFAIP2* expression, changes that would be expected to oppose TNT formation (Brou & Zurzolo, 2026; F. Wang et al., 2021). These changes may represent a compensatory response that limits excessive membrane protrusion following TNT induction, although this possibility requires experimental validation. Lactate enhanced mitochondrial oxidative metabolism in an LDH-dependent but MCT1-independent manner. Thus, although both oxidative metabolism and TNT formation depend on lactate metabolism, these responses appear to diverge downstream of transporter-mediated uptake.

Lactate-treated spheroids were less sensitive to TMZ-induced growth suppression. In addition to the established role of TNT-mediated MGMT transfer in TMZ resistance, tumour microtube networks can protect tumour cells from cytotoxic stress, and mitochondrial transfer from neighbouring cells can restore oxidative metabolism and reduce chemotherapy-induced cell death in other tumour models (Wu et al., 2026). Our findings suggest a related metabolic mechanism in which lactate promotes both oxidative metabolism and TNT-dependent mitochondrial exchange, thereby reducing the response of GBM cells to TMZ. However, we did not determine whether blocking TNT formation restores TMZ sensitivity in this setting. Therefore, a direct link between lactate-induced TNT formation and the spheroid response remains to be established.

Based on these findings, we propose a model in which extracellular lactate is taken up through MCT1 and metabolised by LDH, supporting oxidative phosphorylation and contributing to p300-mediated lactylation. These metabolic and epigenetic signals are associated with increased expression of *CDC42*, *MYO10*, and *FSCN1*, enhanced TNT biogenesis, and mitochondrial transfer. Increased intercellular metabolic connectivity, together with lactate-supported oxidative phosphorylation, may contribute to reduced TMZ sensitivity.

### Limitations

The changes in *CDC42*, *MYO10*, *FSCN1*, *MICAL2PV*, and *TNFAIP2* expression are correlative, and the relative contributions of histone and non-histone lactylation have not been directly established. In addition, the TMZ-response experiments were performed using spheroid models and require validation in orthotopic or patient-derived xenograft models. Despite these limitations, our findings identify lactate metabolism, including MCT1-dependent uptake, LDH-dependent metabolism, and p300-mediated lactylation, as a potential mechanistic link between the GBM metabolic microenvironment, TNT-mediated intercellular communication, and TMZ resistance. This lactate-TNT axis may therefore represent a potential target for therapeutic strategies aimed at limiting tumour cell connectivity during chemotherapy.

## Materials and Methods

### Cell culture and maintenance

Human glioblastoma U-87 MG cells, human breast cancer MDA-MB-231 cells, and HEK293T cells were purchased from NCCS, Pune, India, and cultured in high-glucose Dulbecco’s modified Eagle’s medium (DMEM; Gibco #12100046) supplemented with 10% heat-inactivated foetal bovine serum (FBS; Gibco #A5256701) and 1× Antibiotic-Antimycotic (Anti-Anti; Gibco #15240062). Cells were maintained at 37°C in a humidified atmosphere containing 95% air and 5% CO₂. At 80-90% confluence, cells were subcultured using trypsin solution (Gibco #25300062). For cryopreservation, cells were suspended in medium containing 70% DMEM, 20% FBS, and 10% DMSO.

### Live-cell imaging by fluorescence microscopy

Cells were grown in 35-mm glass-bottom dishes and treated under the indicated conditions for 24 h. Cells were stained with CellMask™ Plasma Membrane Green (Thermo Fisher #C37608) at a final dilution of 1:200 in Hank’s Balanced Salt Solution (HBSS). Images were acquired using an Evident IX83 epifluorescence microscope with a 40X extended-apochromatic objective (0.95 NA). For high-resolution visualisation of TNTs and mitochondria within TNTs, U-87 MG cells were stained with CellMask™ Plasma Membrane Green, MitoTracker™ Red FM (Thermo Fisher #M22425), and Hoechst (Thermo Fisher #H1398). Representative images in Fig. 1A, B were acquired using a FLUOVIEW FV3000 confocal laser-scanning microscope with a 60X oil-immersion objective (X Apo, 1.42 NA).

### TNT identification and quantification

TNTs were visualised at 60-70% cell confluence. The TNT index, defined as the number of TNTs per 100 cells, and TNT length (µm) were quantified manually using ImageJ-Fiji (64-bit) with the Cell Counter plugin. TNTs were identified according to previously reported criteria (Abounit et al., 2015). Briefly, membrane projections had to be suspended above the substratum and connect two distant cells within the middle or upper z-planes.

### Generation of stable cell lines by lentiviral transduction

HEK293T cells were transfected with the packaging plasmid psPAX2 (Addgene #12260), the envelope plasmid pMD2.G (Addgene #12259), and the transfer plasmid pLenti CMV GFP Puro (658-5) (Addgene #17448) or pLV-mitoDsRed (Addgene #44386). Viral supernatant was collected after 48 h and added to U-87 MG cultures in the presence of polybrene. Transduced cells were sorted after 48 h for GFP- or DsRed-positive cells. The stable cell pools were expanded and used for subsequent experiments.

### Mitochondrial transfer assay

Mitochondrial transfer from mito-DsRed-expressing donor cells to GFP-expressing recipient cells was assessed. For co-culture experiments, cells were mixed at a 1:1 ratio and exposed to 10 mM L-lactate for 24 h. For donor preconditioning, mito-DsRed donor cells were treated with 10 mM L-lactate for 24 h and then co-cultured at a 1:1 ratio with untreated GFP-expressing recipient cells for 16 h. Cells were imaged using appropriate filter sets for GFP and DsRed. Mitochondrial transfer was quantified as the percentage of GFP-positive recipient cells containing DsRed-positive puncta. For the extracellular vesicle transfer control, U-87 MG cells expressing mito-DsRed were cultured for 24 h in medium with or without 10 mM lactate. Conditioned media were collected and added to unlabelled U-87 MG cells. Recipient cells were imaged 16 h later.

### 3D spheroid model

U-87 MG-derived spheroids of uniform size were generated by seeding 2,000 cells per well in agarose-coated 96-well plates under the indicated nutrient conditions. All wells contained 10% FBS and 1× Antibiotic-Antimycotic and were maintained at 37°C in a humidified atmosphere containing 5% CO₂. Images were acquired by phase-contrast microscopy using an IX83 inverted microscope with a 10X objective on day 4. Spheroids were then treated with 50 µM TMZ, and images were acquired on day 9. Spheroid diameter was quantified using ImageJ-Fiji. The percentage reduction following TMZ treatment was calculated as: [(equivalent diameter of untreated spheroid − equivalent diameter of TMZ-treated spheroid) / equivalent diameter of untreated spheroid] × 100.

### Real-time qPCR analysis

Total RNA was isolated using RNAiso Plus reagent (Takara #9108) and reverse-transcribed using a reverse-transcription kit (Bio-Rad). Gene expression was measured by quantitative PCR using SYBR Green Master Mix on an Applied Biosystems instrument. Primers were designed using Primer-BLAST (NCBI, NIH) and are listed in Table S1. Relative mRNA expression was calculated using the 2^⁻ΔΔCt^ method with RPL13A as the reference gene. For HCAR1, PCR products were resolved on a 1.5% agarose gel and visualised by ethidium bromide staining.

### Seahorse assay

Oxygen consumption rate (OCR) was measured using an Agilent Seahorse XFe96 Analyzer. Cells (10,000 per well) were seeded in Seahorse XFe96 microplates and cultured under the indicated conditions before extracellular flux analysis. Mitochondrial stress tests were performed by sequential injection of oligomycin (2.5 µM), FCCP (2 µM), and rotenone/antimycin A (0.5 µM). Data were analysed using the Seahorse XF Mito Stress Test Report Generator and normalised to total protein.

### Western blot

Whole-cell lysates were prepared in RIPA buffer, and protein concentrations were determined by Lowry’s method. Protein (50 µg per lane) was separated by 8% SDS-PAGE and transferred to a PVDF membrane. The membrane was blocked with 5% skimmed milk and 0.1% Tween-20 dissolved in Tris-buffered saline (TBS) pH 7.4 for 1 h at room temperature. Following blocking, the membrane was probed with anti-pan lactic acid-lysine primary antibody (Sigma #SAB5701141; 1:1000) overnight at 4 °C followed by HRP-conjugated anti-rabbit secondary antibody (Cell Signaling Technology #7074P2; 1:5000) for 1 h at room temperature. Vinculin was used as a loading control and detected using a mouse anti-vinculin antibody (Santa Cruz Biotechnology #sc-59803; 1:5000), followed by HRP-conjugated anti-mouse secondary antibody (Cell Signaling Technology #7076P2; 1:5000). Protein bands were visualized using Clarity ECL substrate on a Bio-Rad ChemiDoc XRS, quantified by densitometry, and normalised to the corresponding vinculin signal.

### Statistics

Experiments were performed with at least three biological replicates. Data are presented as mean ± SEM unless otherwise stated; spheroid data are presented as mean ± SD. Two conditions were compared using an unpaired t-test or Mann-Whitney U test. Experiments involving more than two conditions were analysed using one-way ANOVA followed by Tukey’s multiple-comparisons test. Spheroid experiments involving multiple conditions and time points were analysed using two-way ANOVA followed by Tukey’s multiple-comparisons test. The statistical test used for each comparison is indicated in the corresponding figure legend. P < 0.05 was considered statistically significant. Analyses were performed using Microsoft Excel and GraphPad Prism. Illustrations were prepared using BioRender.

## CRediT authorship contribution statement

**Sambhavi Pattnaik:** Writing – original draft, Visualization, Validation, Methodology, Investigation, Data curation and analysis. **Subashree Anand and Ganeshini P:** Visualization, Validation, Methodology, Data curation and analysis. **Amal Kanti Bera:** Writing – review & editing, Project administration, Funding acquisition, Conceptualization.

## Declaration of Generative AI and AI-assisted technologies in the writing process

During the preparation of this work the authors used ChatGPT and Claude in order to improve readability and language. After using these tools, the authors reviewed and edited the content as needed and take full responsibility for the content of the published article.

## Funding and Acknowledgements

This work was supported by an Exploratory Research Grant (RF25260612BTRFER008220) from the Indian Institute of Technology Madras. SP was supported by the Prime Minister’s Research Fellowship (PMRF), Ministry of Education, Government of India. The authors acknowledge the BIO-SAIF at the Department of Biotechnology, Indian Institute of Technology, Madras for providing support with the confocal microscope and the Agilent Seahorse XF Analyzer.

## Data availability

Additional data are provided in the Supplementary Materials.

