## supplementary_data for "Lactate Promotes Tunneling Nanotube Formation and Reduces Temozolomide Sensitivity in Glioblastoma Cells"

**Fig.S1**

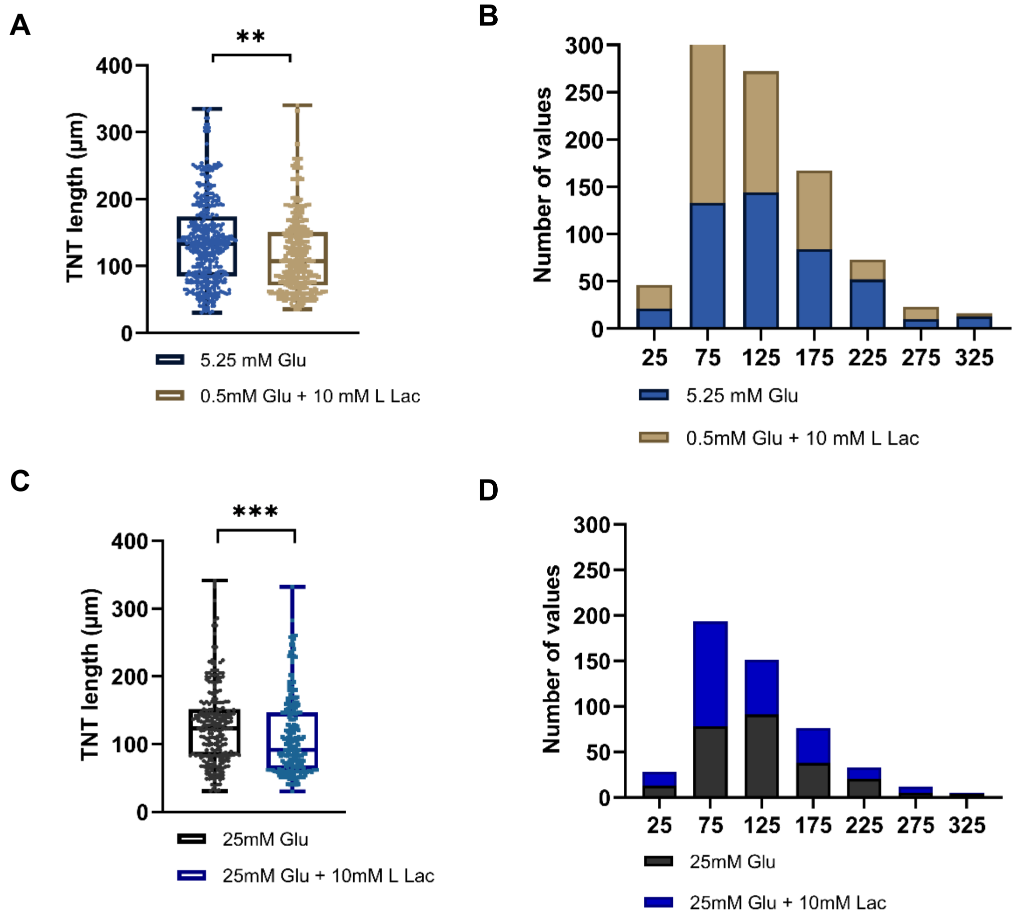


**Fig.S1 Extracellular lactate shifts the TNT length distribution towards shorter TNTs**

***(A)*** *TNT length in U-87 MG cells cultured in 5.25 mM glucose or 0.5 mM glucose + 10 mM L-lactate (n = 450 TNTs). Each point is one TNT; boxes show the median and interquartile range, whiskers the minimum and maximum. **P < 0.01, Mann-Whitney U test.* ***(B)*** *Frequency distribution (number of TNTs) of the TNT lengths in (A), in 50-µm bins.* ***(C)*** *TNT length in cells cultured in 25 mM glucose or 25 mM glucose + 10 mM L-lactate (n = 250 TNTs). ***P < 0.001, Mann-Whitney U test.* ***(D)*** *Frequency distribution (number of TNTs) of the TNT lengths in (C), in 50-µm bins.*

**Fig.S2**

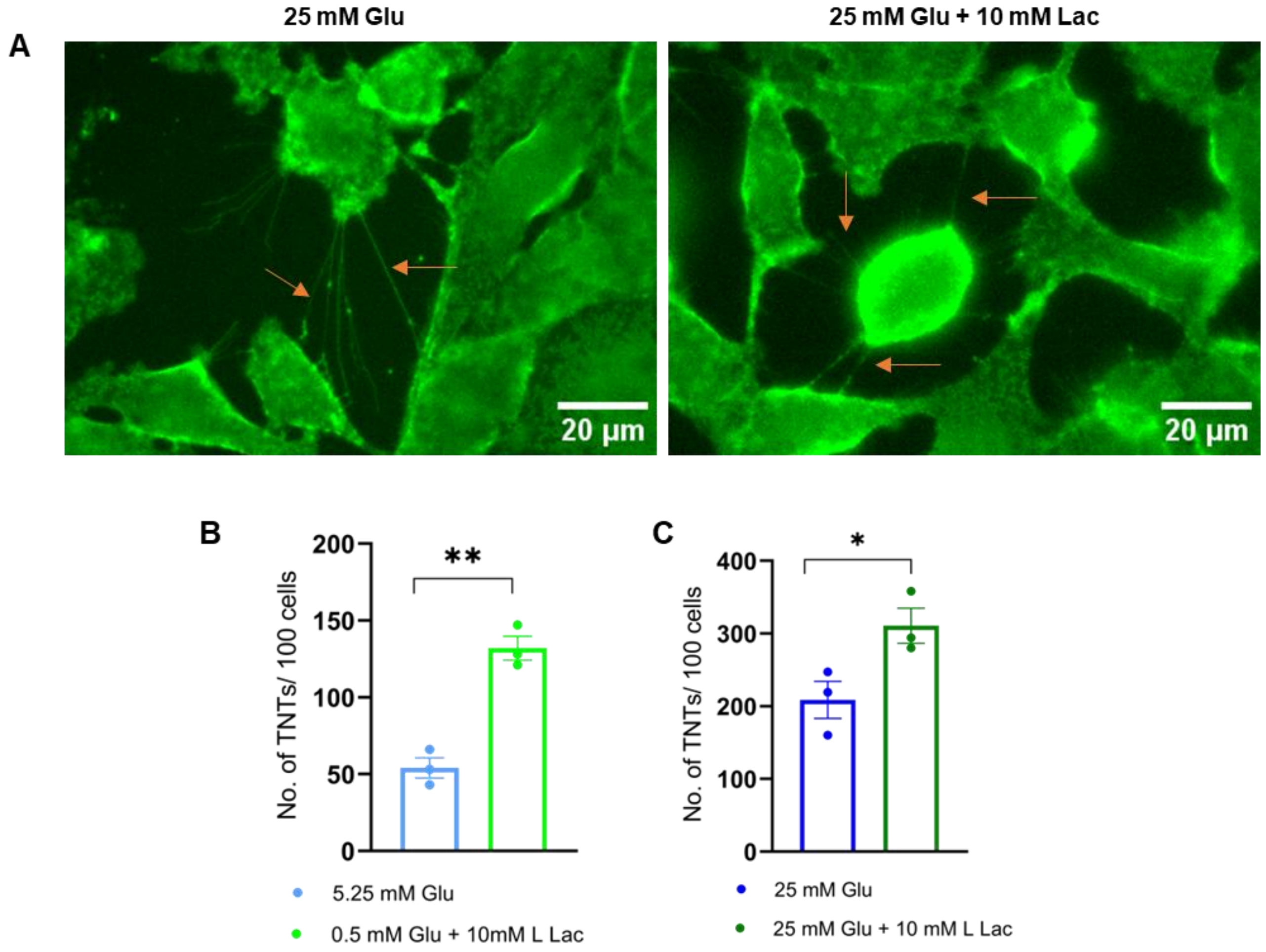


**Fig. S2 Lactate increases TNT formation in MDA-MB-231 breast cancer cells**

*(A) Representative images of MDA-MB-231 cells cultured in 25 mM glucose or 25 mM glucose + 10 mM L-lactate and stained with CellMask™ Plasma Membrane Green.* ***(B)*** *TNT number per 100 cells in 5.25 mM glucose vs 0.5 mM glucose + 10 mM L-lactate. **$P < 0.01$, unpaired t-test.* ***(C)*** *TNT number per 100 cells in 25 mM glucose vs 25 mM glucose + 10 mM L-lactate. *$P < 0.05$, unpaired t-test. Data are mean ± SEM from three independent experiments. Scale bar, 20 μm.*

**Fig.S3**

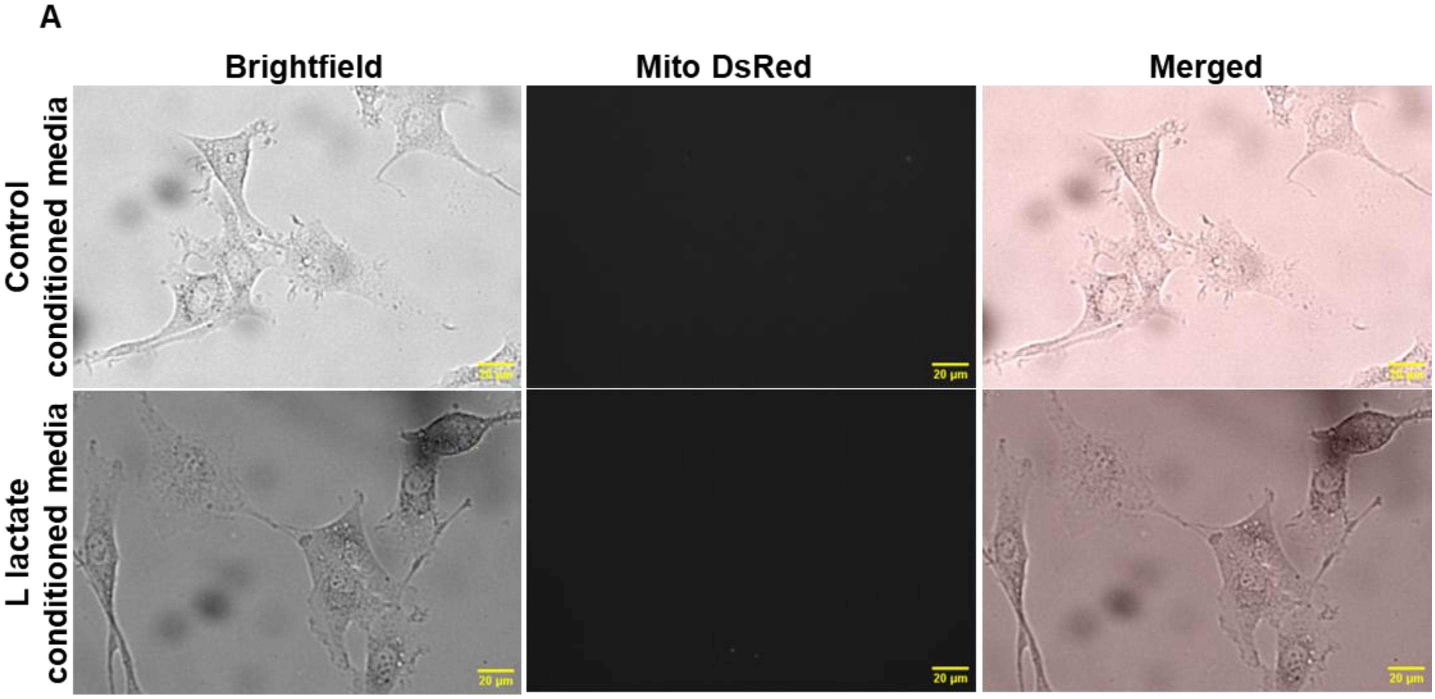


**Fig.S3 Lactate does not enhance extracellular vesicle (EV)-mediated mitochondrial transfer**

***(A)*** *Representative brightfield, Mito-DsRed, and merged images of recipient U-87 MG cells 16 h after incubation with conditioned media collected from U-87 MG cells expressing DsRed-labelled mitochondria, cultured for 24 h in the absence or presence of 10 mM L-lactate. Negligible DsRed-positive puncta are visible in recipient cells under both conditions, with no apparent difference between the two groups, indicating that EV-mediated mitochondrial transfer is not increased by lactate treatment and contributes minimally to the mitochondrial transfer observed via TNTs. Scale bar, 20 µm.*

**Fig.S4**

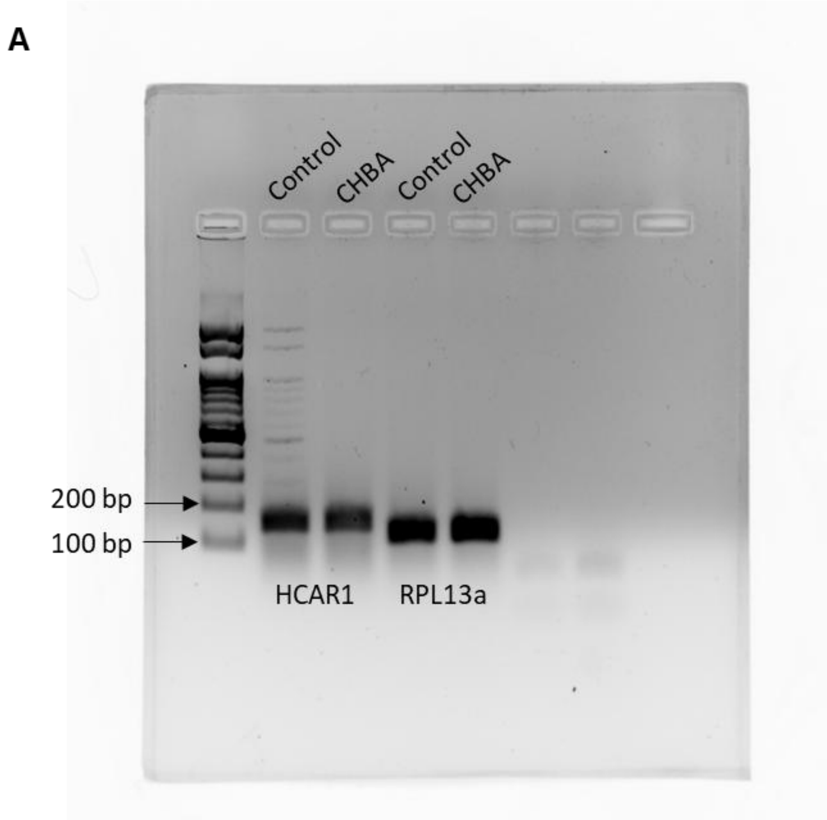


**Fig.S4 Semi-quantitative PCR validation of HCAR1 expression**

***(A)*** *Agarose gel electrophoresis of PCR products amplified from U-87 MG GBM cells treated with CHBA or vehicle control. HCAR1 transcript levels were assessed relative to the housekeeping gene RPL13a.*

**Table S1: Primers for RT-PCR**

| Gene | Forward Primer Sequence | Reverse Primer Sequence |
|---|---|---|
| *TNFAIP2* | TGCCTGTCCTTCCGGATGTC | GTGGCTCTTGAGCTCACCCA |
| *MYO10* | GCACTATGCTGGAGAGGTGCAA | CGCCGATGTTTGCTTCCACA |
| *CDC42* | TGTTCTGCACTTACACAGAAAGGC | CATAGCAGCACACACCTGCG |
| *FSCN1* | GTGGAGACAGCAGGGGACTC | GACGTCATAGCTGGAGCGG |
| *MICAL2PV* | CCAGGTATCTCCACCTCCTT | AGTCCCTGCATCCAGTTACAC |
| *RPL13A* | CCTGGAGGAGAAGAGGAAAGAGA | TTGAGGACCTCTGTGTATTTGTCAA |
